# Filamentation and inhibition of prokaryotic CTP synthase

**DOI:** 10.1101/2023.10.19.563106

**Authors:** Chen-Jun Guo, Zi-Xuan Wang, Ji-Long Liu

**Affiliations:** School of Life Science and Technology, ShanghaiTech University, Shanghai, China; Department of Physiology, Anatomy and Genetics, University of Oxford, Oxford, UK

**Keywords:** CTP synthase, cytoophidium, metabolic filament, Cryo-EM, species specific inhibition

## Abstract

CTP synthase (CTPS) plays a pivotal role in the de novo synthesis of CTP, a fundamental building block for RNA and DNA, which is essential for life. CTPS is capable of directly binding to all four nucleotide triphosphates: ATP, UTP, CTP, and GTP. Furthermore, CTPS can form cytoophidia in vivo and metabolic filaments in vitro, undergoing regulation at multiple levels. CTPS is considered a potential therapeutic target for combating invasions or infections by virus or prokaryotic pathogens. Utilizing cryo-electron microscopy, we have determined the structure of *Escherichia coli* CTPS (ecCTPS) filament in complex with CTP, NADH, and the covalent inhibitor DON, achieving a resolution of 2.8Å. We construct a phylogenetic tree based on differences in filament-forming interfaces and design a variant to validate our hypothesis, providing an evolutionary perspective on the CTPS filament formation. Our computational analysis reveals a solvent-accessible ammonia tunnel upon DON binding. By comparative structural analysis, we discern a distinct mode of CTP binding of ecCTPS, differing from eukaryotic counterparts. Combining biochemical assays and structural analysis, we determine and validate the synergistic inhibitory effects of CTP with NADH or adenine on CTPS. Our results expand our comprehension of diverse regulatory aspects of CTPS and lay a foundation for the design of specific inhibitors targeting prokaryotic CTPS.

## Introduction

CTP synthase (CTPS) plays a critical role in catalyzing the final and rate-limiting step of de novo CTP synthesis. CTPS is composed of a glutamine amidotransferase (GAT) domain and a kinase-like ammonia ligase (AL) domain[1, 2]. It converts the substrate UTP into CTP, utilizing ammonia generated from glutamine and ATP as an energy source[3]. Considering the indispensability of its product CTP in DNA, RNA and phospholipid synthesis, CTPS is meticulously regulated across various layers [4–11]. While CTP can inhibit its activity, GTP serves as an enzymatic regulator, stimulating the reaction at low concentrations and inhibiting it at higher levels. The redox cofactor NADH has been found to inhibit ecCTPS, holding potential physiological relevance [12].

In solution, *Escherichia coli* CTPS (ecCTPS) exists as an inactive dimeric form under dilute conditions. Upon the addition of substrate UTP and ATP, it transitions into an active tetrameric form, which is also observed in eukaryotic CTPS[13]. The presence of product CTP prompts CTPS to shift into an inactive tetrameric state for both prokaryotic and eukaryotic[14]. Recent studies have revealed that CTP can bind to CTPS at two distinct binding sites for various eukaryotic species[15–17].

Beyond oligomerization, CTPS demonstrates the capability to further aggregate, forming metabolic filaments[15, 18–22]. Filaments of CTPS have been confirmed and characterized in multiple species and conditions, where ecCTPS has been shown to form large-scale, inhibitory filaments in the presence of product CTP[19]. Both human hCTPS1 and hCTPS2, along with Drosophila CTPS, can form different filaments under the conditions of ATP, UTP, or CTP, with varying effects on reaction promotion. Ura7 and ura8 from budding yeast also form filaments under substrate and product conditions and respond to changes in pH[15].

CTPS has been observed to aggregate into micron-scale non-membrane-bound organelles named cytoophidium (plural cytoophidia), due to their snake-like appearance[18, 23, 24]. Cytoophidia are highly conserved across multiple species and all three domains of life, suggesting their special biological roles. In prokaryotes, cytoophidia were initially observed and demonstrated to be crucial for the morphology of bacteria, such as *Caulobacter crescentus*[18]. *Drosophila* have served as a classic model for studying CTPS cytoophidia[25]. Through experiments involving point mutations and others, CTPS cytoophidia have been found to have close ties to adipose tissue architecture[26].

Despite its complex regulation, the vital biological role of CTPS positions it as a potential drug target for diseases such as cancer[27], bacterial infections[28], and parasitic infections[29]. As our understanding deepens, the design for targeted regulation of CTPS and species-specific regulation are becoming feasible. Recently, Matthew J. McLeod et al unveiled the inhibitory effects of gemcitabine-5′-triphosphate, a therapeutic agent against solid tumors, on ecCTPS. It exhibited strong binding affinity, approximately 80-fold greater than CTP, providing novel insights into inhibitor development[30]. Utilizing cryo-EM, Eric M. Lynch et al revealing the mechanisms of selective inhibition of human CTPS and lay a foundation for the design of immunosuppressive therapies[16].

In this study, we solve the structure of ecCTPS in complex with CTP, NADH, and the covalent inhibitor DON. Incorporating phylogenetic analysis and bioengineering, we gain evolutionary insights into CTPS filament formation. Through computational analysis, we reveal a solvent-accessible ammonia tunnel upon DON binding. Our structures capture the interaction between NADH and ecCTPS, and biochemical assays reveal that NADH achieves synergistic inhibition by interacting with the ATP binding pocket through its Adenine moiety. Furthermore, we identify distinct CTP binding and inhibition modes in ecCTPS compared to already resolved eukaryotic CTPS structures due to sequence variations. Our findings potentially lay the groundwork for developing species-specific CTPS inhibitors.

## Result

### Structure of ecCTPS bound with CTP, NADH and DON

Recently, it has been discovered that CTPS possesses the intriguing ability to spontaneously assemble into filaments in vitro under different conditions. Structures of CTPS filament have been solved in various eukaryotic organisms, including human, drosophila, and budding yeast. Previously, researchers have observed and characterized CTPS filaments from the prokaryote in vitro and in vivo.

In pursuit of establishing a more robust structural foundation for the study of prokaryotic CTPS filaments, we employed a recombinant expression approach to purify and isolate the CTPS protein from *Escherichia coli* (Fig S1). Utilizing cryo-electron microscopy (cryo-EM) and single particle analysis, we resolved the structure of purified ecCTPS bounded with CTP, NADH and 6-diazo-5-oxo-norleucine (DON), achieving a resolution of 2.8 Å (Fig 1A) (Fig S2-4). During the sample preparation process, under conditions with product CTP, we observed a distinct filamentous arrangement of ecCTPS in the cryo samples (Fig S3). The results of 2D classification provided a clear visualization of ecCTPS filament, wherein it assembles into filaments with CTPS tetramers as the helical unit (Fig S3).

**Figure 1.**
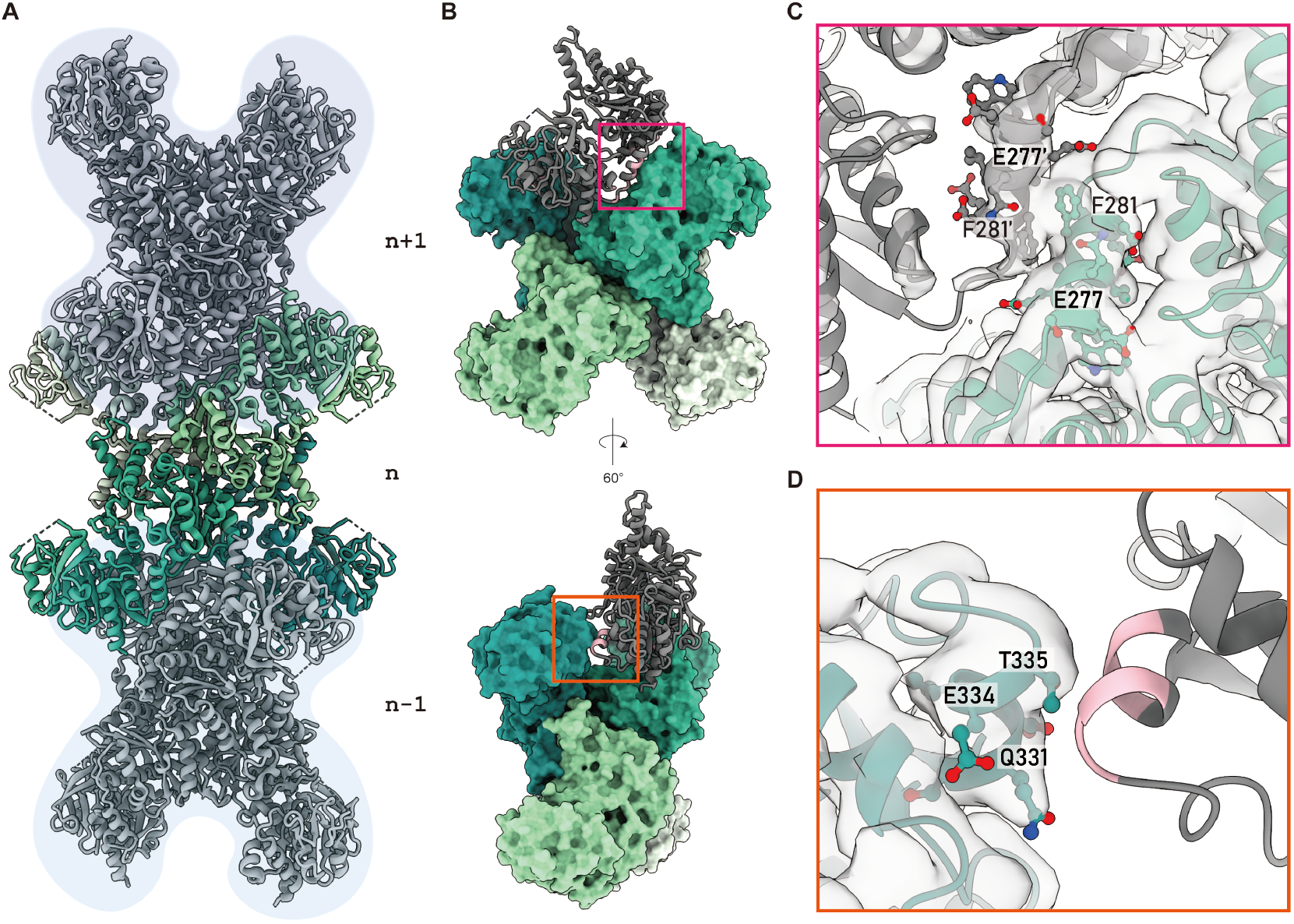
Assembly of ecCTPS filament. **(A)** Structure of the ecCTPS filament. The model is composed of three helical units. The central ecCTPS tetramer is colored by different protomers. The upper and lower layers of the tetramer are depicted in gray ribbon with a light blue background. (**B)** Contact Sites between adjacent Tetramers. One ecCTPS tetramer is shown by the solvent-accessible surface, with another protomer adjacent to ecCTPS depicted by a gray ribbon. Amino acids within 4.5 Å distance are highlighted in pink. Two different angled views are presented. **(C, D)** Major assembly interfaces of the ecCTPS Filament. Zoomed-in views of the magenta and orange boxes in Figure 1B. The map density is displayed as a transparent white surface.

Consistent with findings previously elucidated by Rachael M Barry et al, the interface of the ecCTPS filament comprises two major interaction regions (Fig 1B). One of these regions is located at the linker domain of CTPS, where chains A and A’ engage in a symmetric complementary way (Fig 1C). Notably, the density of residue F281 appears to participate in a direct touch with the density of F281’. These interactions likely contribute to the stability of the ecCTPS filament. The second interface is situated between helices 330-335 within the GAT domain (Fig 1D).

### The evolution of CTPS filament assembly

In comparison to eukaryotic CTPS, the assembly way of ecCTPS filament exhibits significant distinctions. The assembly interface of solved eukaryotic CTPS filaments is primarily centered on helix 12, a sequence that is absent in the E. coli counterpart (Fig S5). In order to systematically analyze this phenomenon from an evolutionary perspective, we selected CTPS sequences from 112 species for the construction of a phylogenetic tree. This dataset included 50 prokaryotes, 23 eukaryotes, and 39 archaea. The presence or absence of α helix 12 was used as a classification criterion (Fig 2A) (Fig S6).

**Figure 2.**
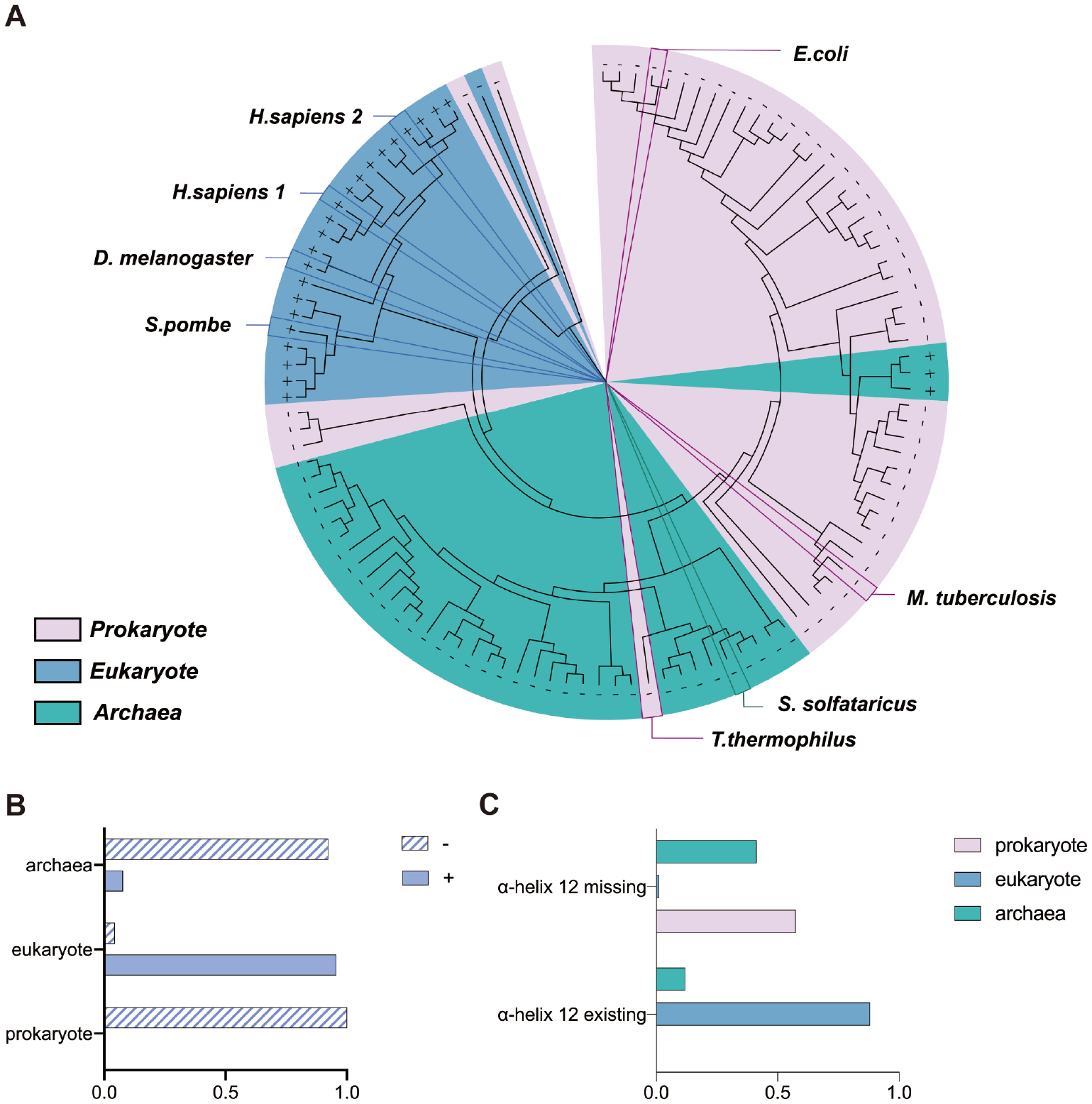
Phylogenetic analysis of CTPS. **(A)** Phylogenetic tree of CTPS consisting of 112 CTPS sequences. “+” represents the existence of α helix 12, and “-” represents the missing of α helix 12. Eukaryotes, prokaryotes, and archaea are distinguished by different colors. **(B)** Classification by species. This histogram indicates the ratios of the existence or missing of α helix 12 in eukaryotes, prokaryotes and archaea. **(C)** Classification by the existence or missing of α helix 12. The ratios of eukaryotes, prokaryotes and archaea in the α helix 12 existing and missing conditions are shown, respectively.

The evolutionary analysis reveals that among the selected 112 species, α helix 12 is present in nearly all eukaryotes (>95%), while it is absent in all prokaryotes and the majority of archaea (88%) (Fig 2B). Based on the presence or absence of α helix 12 for sequence classification, we observed that the majority of organisms possessing helix 12 are eukaryotes, with only a small fraction belonging to archaea. Conversely, among the species lacking α helix 12, prokaryotes and archaea constitute the predominant groups. Notably, within the analyzed dataset, only one eukaryotic organism is devoid of α helix 12 (Fig 2C).

### Insertion of helix12 allows ecCTPS to form filaments with substrates

Based on both previous research and our experimental findings, it is established that the wild-type ecCTPS is capable of forming filaments in the presence of the product CTP. However, filament formation was not observed in the presence of substrates UTP or ATP.

In order to elucidate the necessity and role of helix 12 in CTPS filament formation, we engineered, expressed, and purified ecCTPS with an inserted hCTPS helix 12 (referred as ecCTPS^helix12+^)(Fig S7).

We employed AlphaFold2 to predict the structure of ecCTPS^helix12+^. Among the five structures predicted, rank 2 exhibited the best performance across various criteria, leading us to select this model for subsequent analysis (Fig S8). Structural alignment revealed that ecCTPS^helix12+^ possesses a markedly different secondary structure compared to wild type ecCTPS. The insertion of the human CTPS sequence resulted in the extension of the originally shorter loop337-341, forming a complete helix that bears a resemblance to α helix12, responsible for filament assembly in eukaryotic CTPS(Fig 3A). When we aligned the predicted model with hCTPS1 filament based on the filament assembly interface, we observed a high degree of similarity, with an RMSD of 0.479 angstroms. H346 and W349 exhibited patterns consistent with His355 and Trp358 of hCTPS1, which play vital roles in the filament assembly(Fig 3B). This implies that ecCTPS^helix12+^ possesses a structural basis for assembling into filaments resembling those in hCTPS1.

**Figure 3.**
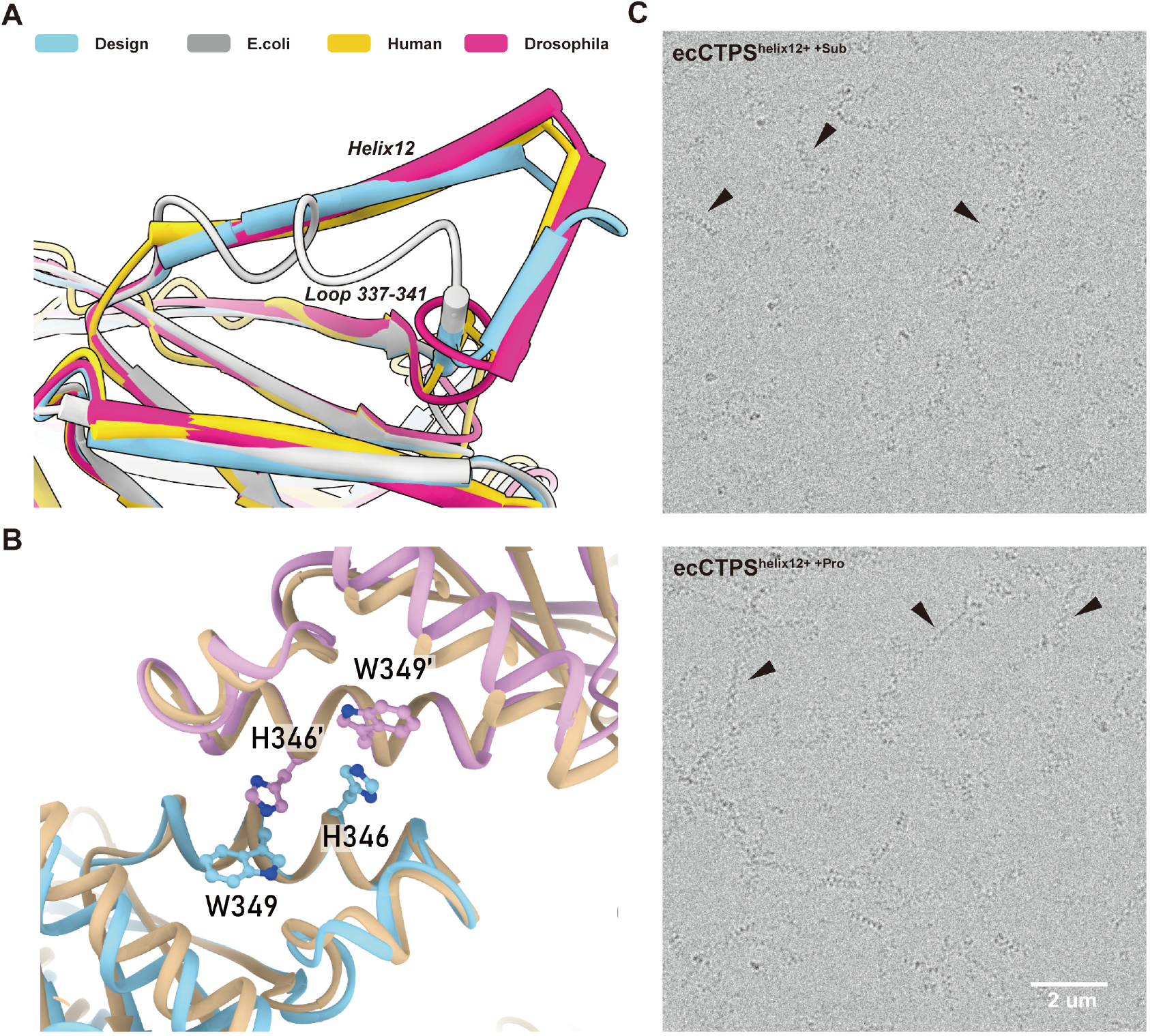
Properties of ecCTPS^helix12+^. **(A)** Comparison of helical assembly interface with predict structure of ecCTPS^helix12+^. The structure of design is predicted by the AlphaFold2. Models are colored according to the color key. **(B)** Simulated helical interface of ecCTPS^helix12+^. Two monomers of ecCTPS^helix12+^ are aligned to the structure of hCTPS1 filament by helix12. **(C)** Cryo-electron micrograph of ecCTPS^helix12+^ under substrate and product conditions. Filaments are indicated by black arrows. Scale bar is defined as 2μm.

To further investigate the nature of ecCTPS^helix12+^, we prepared samples of it in both substrate and product states. Upon analyzing cryo-samples of ecCTPS^helix12+^, we observed that this engineered variant not only formed filaments in the presence of the product CTP but also exhibited filament formation in the presence of substrates UTP and ATP (Fig 3C). Intriguingly, the arrangement of the formed filaments exhibited a connected X-shaped pattern, resembling the previously observed helix 12-mediated helical interface in eukaryotic CTPS filaments. This result demonstrates that the insertion of hCTPS helix 12 is sufficient to alter the assembly mode of ecCTPS filaments, suggesting the pivotal role of helix 12 in determining the CTPS filament assembly pattern.

Our study establishes a connection between protein sequences and the assembly of CTPS filaments, while also providing an evolutionary perspective on the formation of CTPS filaments.

### DON binding and ammonia tunnel formation of ecCTPS

CTPS is composed of GAT and AL domains, and during the reaction, the intermediate product ammonia is transferred between the two domains through an internal gas tunnel. DON is an irreversible inhibitor of CTPS, which can covalently bind to CTPS and inhibit its activity.

In 1971, A. Levitzki et al. demonstrated that even after DON binding, prokaryotic CTPS can still use NH4+ as a substrate for CTP synthesis[31]. In our obtained structure, DON forms a covalent bond with C359, a residue critical for the catalytic function of ecCTPS. Surrounding amino acids, such as F353, contribute to its binding and stabilization (Fig 4A). Upon the conjunction of DON, the GAT domain adopts a closed conformation, preventing external NH4+ from accessing the ammonia tunnel through the catalytic pocket of GAT for further utilization by the AL domain.

**Figure 4.**
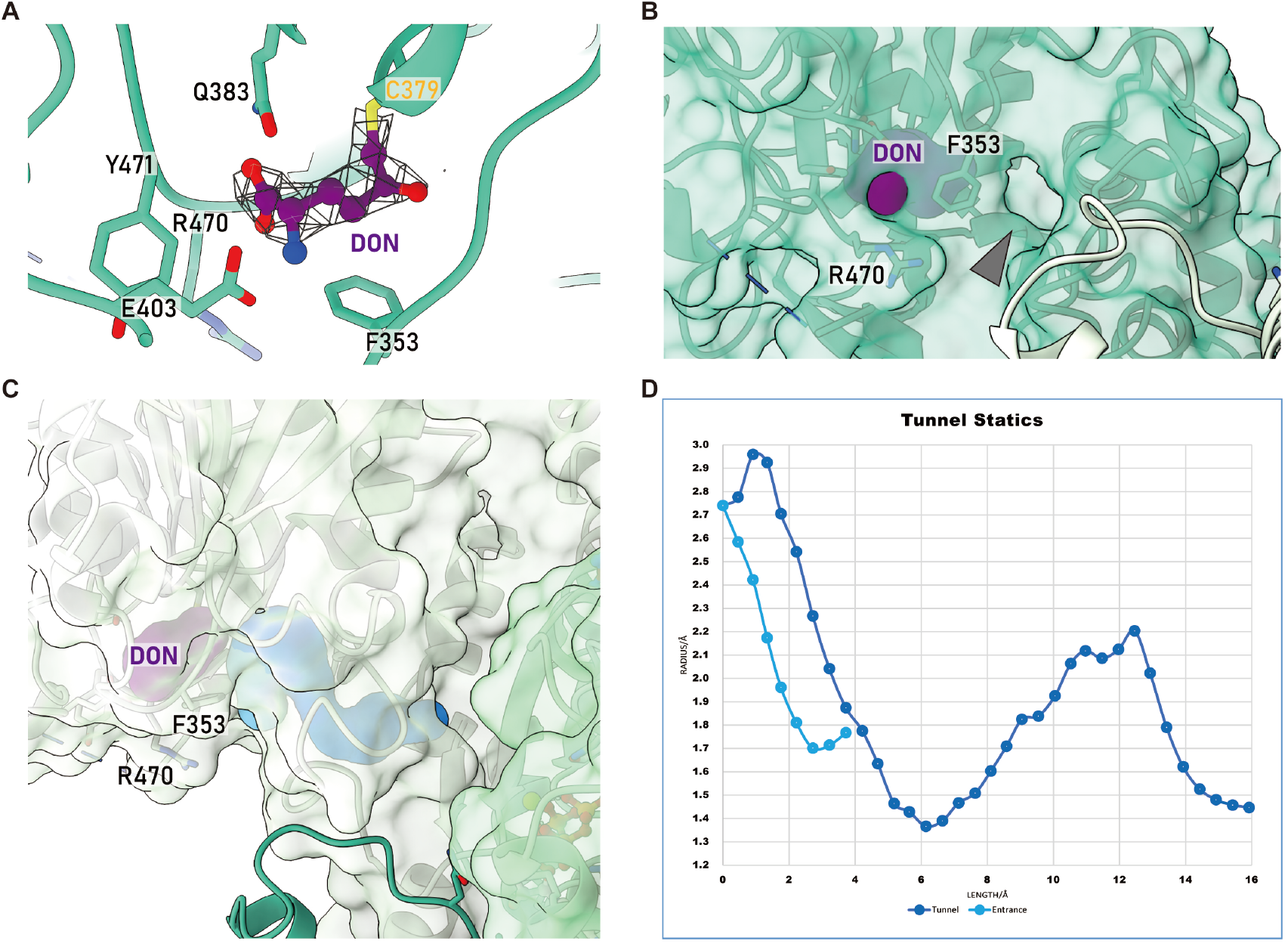
Ammonia tunnel of DON inhibited ecCTPS. **(A)** Binding of DON to ecCTPS. DON is represented by the purple ball-and-stick model, and the density map is depicted as a mesh. **(B)** Ammonia entrance during DON inhibition. The surface of ecCTPS is depicted in transparent green, while the surface of DON is shown in purple. Gray arrows indicate the location of the entrance. **(C)** Ammonia tunnel of DON-inhibited ecCTPS. The surface of ecCTPS is transparent, with the entrance and ammonia tunnel represented in light blue and deep blue, respectively. **(D)** Tunnel statics. The two curves in light blue and deep blue correspond to the parameter of Entrance and Tunnel in Figure 3C. The x-axis represents the distance along the channel center, while the y-axis represents the maximum accessible radius from the corresponding channel center, measured in angstroms.

Through computational analysis, we observed the presence of a solvent-accessible channel on the side of F353 close to the AL domain, connecting an internal cavity within the GAT domain (Fig 4B and C). The minimum diameter of this channel exceeds 1.7 Å allowing the passage of NH4+ (Fig 4D). This cavity is also positioned above the ammonia tunnel that connects to the AL domain, where we identified a minimum diameter of the ammonia tunnel above 1.3 Å, allowing the transfer of the intermediate products. The existence of this entrance of ammonia tunnel explains the feasibility of ecCTPS utilizing NH4+ as a substrate when DON bound to the GAT pocket.

### Diverse binding modes of CTP in CTPS

The product CTP serves as a natural inhibitor of various eukaryotic CTPS and can bind to the substrate binding pockets of UTP or ATP in two distinct inhibitory modes: competitive and non-competitive with substrate UTP.

In our structure of ecCTPS, CTP binding was observed in the UTP-binding pocket(Fig 5A), consistent with previously resolved ecCTPS-CTP complexes (Fig 5B). CTP shares the triphosphate binding site with UTP, and interactions between coenzyme magnesium ions and the triphosphate moiety further stabilize CTP binding. The cytidine head and ribose of CTP are enveloped by three adjacent protomers, stabilized through electronic interactions(Fig 5A&B).

**Figure 5.**
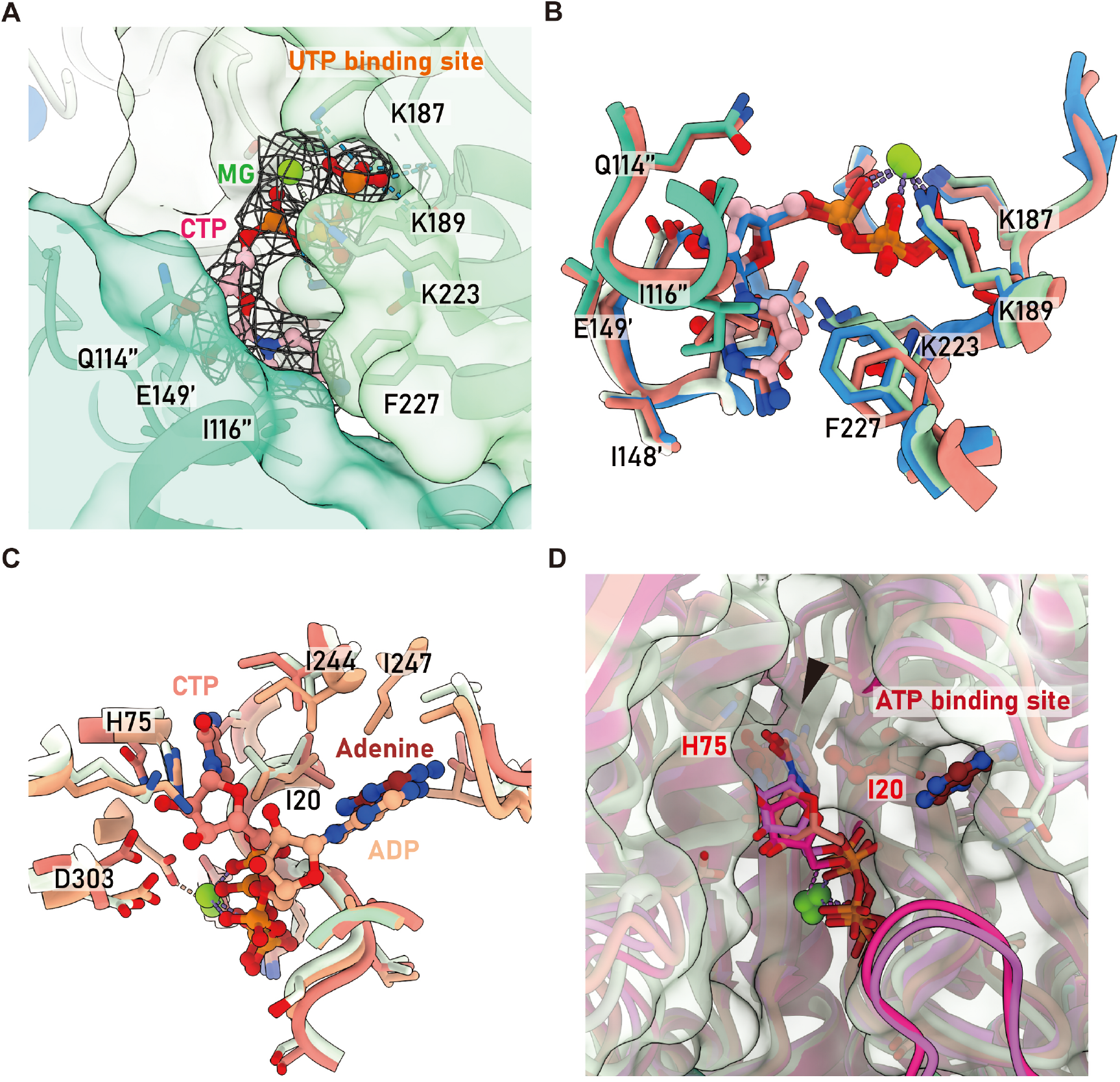
Comparisons of CTP binding sites. **(A)** Classical CTP binding pocket. CTP is represented by pink sphere- and-stick models, ecCTPS is color-coded by protomers, map density is depicted as a mesh, and hydrogen bonds are shown as blue dashed lines. **(B)** Comparisons of classical CTP binding site. Models of dmCTPS, ecCTPS (X-Ray), and ecCTPS (EM) are colored in reddish, blue, and green, respectively, with corresponding PDB codes (7DPW, 5TKV, 8I9O). **(C)** Comparisons of non-classical CTP binding site. Models of dmCTPS with CTP binding in both classical and non-classical pockets, ecCTPS with ADP, CTP binding, and the model of this study are displayed, with PDB codes (7DPW, 2AD5, 8I9O). They are colored in reddish, beige, and green, respectively. **(D)** Comparison of allosteric inhibitory sites for CTP binding. The clash of non-classical CTP binding with the simulated solvent-accessible surface of ecCTPS is indicated by the black arrow. Pink, reddish, and magenta ribbons respectively represent models of dmCTPS, hCTPS1, and hCTPS2, PDB codes (7DPW, 7MH0 and 7MH1). Structures are aligned by fitting into the map of ecCTPS.

Previous research has revealed that the product CTP can bind to CTPS near the substrate ATP binding pocket. This binding mode, initially discovered in dmCTPS in 2021, emerged years after the classical CTP binding site was identified. As a result, this unique CTP binding mode was coined as the ‘non-classical binding of CTP.’ Subsequently, the existence of non-classical CTP binding mode was also confirmed in both hCTPS1 and hCTPS2.

In this non-classical binding mode, the triphosphate moiety of CTP shares the same pocket as ATP, while the cytidine head of CTP adopts a distinct binding mode alongside the adenine head of ATP(Fig 5C). However, in proximity to the ATP-binding pocket, we did not observe the density of CTP in our map. Aligning the structure with those of dmCTPS and hCTPS containing non-canonical CTP binding, we identified a clash between the CTP head of the eukaryotic CTPS model and the solvent-accessible surface of ecCTPS model due to the hindrance by amino acids His75 and Ile20 (Fig 5D). This clash impedes CTP from binding to ecCTPS in a non-competitive inhibitory mode similar to that in eukaryotic CTPS, within the ATP pocket.

### NADH binds to ecCTPS in the ATP binding pocket

In earlier research, C. Habrian et al. discovered that NADH can synergistically inhibit CTPS along with CTP[12]. However, the molecular basis for this regulation is missing. We observed NADH binding in the ATP-binding pocket in our model (Fig 5D). At the 1σ counter level, the adenine head of NADH exhibited well-defined density, while the remaining portion lacked stable density (Fig 6A).

**Figure 6.**
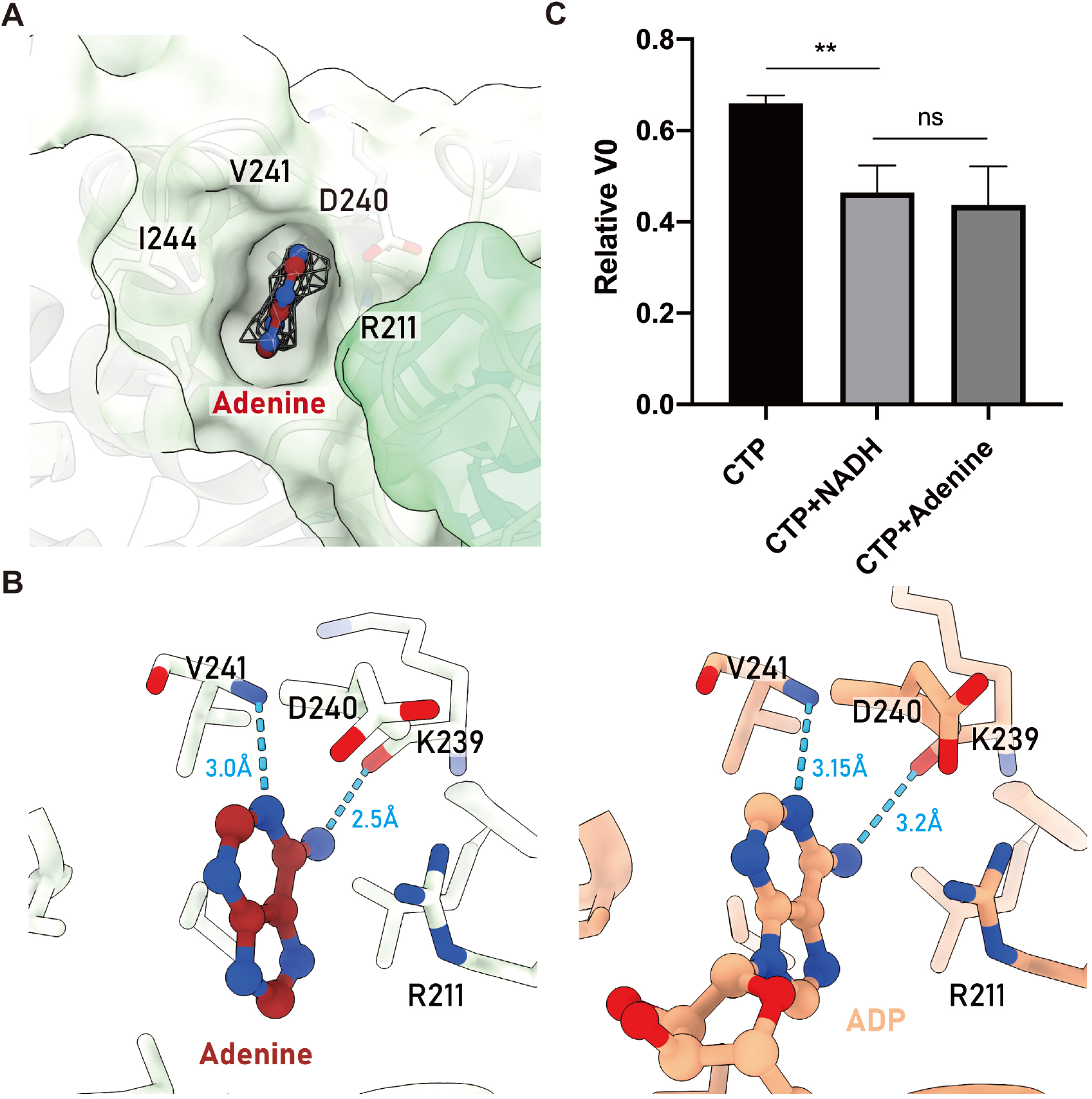
NADH binding and inhibition. **(A)** Density of the adenine moiety of NADH. Map density is depicted as a mesh. **(B)** Comparison of NADH and ADP binding. Hydrogen bonds are indicated by the blue dashes. Left: Model of this study. Right: PDB 2AD5. **(C)** Enzyme activity assays for inhibition by CTP, NADH, and Adenine on ecCTPS. The initial velocity (V0) in the absence of inhibitor is the benchmark. The relative V0 in the presence of CTP, CTP and NADH, and CTP and adenine are 0.66, 0.46 and 0.44 respectively.

Within the ATP pocket, the adenine moiety of NADH exhibits a mode of interaction similar to that of ADP with the protein. The N1 atom forms a set of hydrogen bonds with the nitrogen atom on the V241 backbone, with a distance of 3 angstroms. Additionally, the N6 atom forms another set of stronger hydrogen bonds with the oxygen atom on the K239 backbone, with a distance of 2.5 angstroms. Furthermore, adenine is within 3.8 angstroms of the plane of the R211 side chain, forming a strong pi-pi interaction, which, along with surrounding hydrophobic amino acids, stabilizes the binding of adenine (Fig 6B). According to our model, the adenine moiety of NADH exhibits a stronger interaction with the ATP binding pocket of ecCTPS compared to the adenine moiety of ADP (Fig 6B). This could be attributed to the NADH’s remaining functional group failing to achieve stable binding, thereby providing adenine with ample space for optimization in its binding. This also suggests that NADH’s interaction with ecCTPS may depend on the adenine head.

### Synergistic inhibition of ecCTPS by CTP with NADH or Adenine

To validate the binding effect of NADH and its regulatory role on ecCTPS, we examined the inhibitory effects of both NADH and adenine in combination with CTP on CTPS activity (Fig 6C). As control, CTP alone significantly inhibited CTPS activity. The additional presence of NADH further suppressed the reaction, and the combination of adenine and CTP exhibited a similar degree of inhibition. Enzyme activity assays confirmed the synergistic inhibition of CTPS activity by NADH and CTP and indicated that adenine also possesses this capability.

Our structure provides a structural foundation for the synergistic inhibition of ecCTPS by NADH and CTP. Furthermore, through enzyme activity assays, we propose that NADH’s inhibition of CTPS is likely dependent on the affinity between the adenine head and the ATP pocket of ecCTPS.

## Discussion

In this study, we have determined the structure of complex formed by ecCTPS with CTP, DON, and NADH, shedding light on the interaction pattern between CTPS and the novel ligand NADH. Moreover, our findings refine our comprehension of the regulation exerted by classical ligands like CTP and DON on CTPS. The near-atomic resolution structures provide robust support and validation for our conclusions.

Despite these advances, there are some limitations in our research stemming from experimental constraints. We based our criteria for evolutionary analysis solely on the presence or absence of alpha helix 12. However, it is important to note that there are other differences that could potentially influence filament formation, such as the non-conserved alpha helix 18 at the C-terminal, which appears to have a possible connection to evolutionary relationships. Furthermore, the engineered ecCTPS, successfully assembled into filaments with substrates, displayed certain morphological similarities to eukaryotes. Further investigation is required to determine whether the assembly mode and the regulation pattern for ecCTPS are identical.

Nevertheless, our phylogenetic analysis and bioengineering offer valuable insights. Specifically, it raises intriguing questions about how the two types of filaments, under the product state and substrate state, respectively, evolved. Our findings may imply an asynchronous evolution, prompting us to delve deeper into the significance of these distinct filamentation events in vivo.

## Data availability

The structure data accession codes are EMD-35278 and PDB-8I9O.

## Acknowledgments

EM data were collected at the ShanghaiTech Cryo-EM Imaging Facility. We thank the Molecular and Cell Biology Core Facility (MCBCF) at the School of Life Science and Technology, ShanghaiTech University and Shanghai Frontiers Science Center for Biomacromolecules and Precision Medicine for providing technical support. This work was supported by grants from: Ministry of Science and Technology of China (No. 2021YFA0804700), National Natural Science Foundation of China (No. 31771490), Shanghai Science and Technology Commission (No. 20JC1410500), UK Medical Research Council (grant nos. MC_UU_12021/3 and MC_U137788471) for grants to J.L.L.

## Author contributions

C.J.G initiated the project, developed the protein expression and purification procedures, prepared protein samples for EM studies, collected cryo-EM data, performed data processing and analysis, model building, structure refinement and analyzed experiments. Z.X.W expressed and purified the proteins, helped the samples preparations for EM studies, examined the EM samples, designed and purified the engineered proteins, performed the functional assays and the phylogenetic analysis. C.J.G and Z.X.W visualized results and wrote the manuscript. J.L.L revised manuscript, obtained the funding and provided the experimental resources.

