## Supplemental Info for "Filamentation and inhibition of prokaryotic CTP synthase"

**Supplemental Information**

- **Methods**
- **Supplemental Table S1**
- **Supplemental Figures S1-S8**

### Methods

#### *ecCTPS plasmid construction and protein purification*

The full length ecCTPS gene was cloned into pET28a vector with a C-terminal 6 × His tag and transformed into Escherichia coli Transetta (DE3) cells for expression. Transformed cells were cultured in LB medium at 37 °C with 220 rpm until OD600 reached a range of 0.6 to 0.8, and then induced with 0.1mM IPTG for 16-18 hr at 16 °C. Cells were pelleted by centrifugation at 4,500 rpm for 10 min followed by resuspension in cold lysis buffer (50 mM Tris-HCl pH 7.5, 500 mM NaCl, 10% glycerol, 20mM imidazole, 1 mM PMSF, 5mM β-mercaptoethanol, 5mM benzamidine, 2 μg/ml leupeptin, and 2 μg/ml pepstatin). Bacteria in lysis buffer were disrupted by high pressure at 800 bar and centrifuged at 16,000 × g for 30 min at 4 °C to collect supernatant. This was incubated with equilibrated Ni-Agarose (Qiagen) for 1.5 hr. Next, Ni-Agarose was washed by washing buffer (50 mM Tris-HCl pH 7.5, 500 mM NaCl, 10% glycerol, 40mM imidazole, 5 mM β-mercaptoethanol), and proteins were eluted with elution buffer (30 mM Tris-HCl pH 7.5, 100 mM NaCl, 240 mM imidazole). SuperoseTM 6 Increase 10/30 GL column and AKTA Pure (Cytiva) were used for further purification of peak fractions. Finally, CTPS was eluted with buffer containing 100 mM NaCl and 30 mM Tris-HCl pH 7.5.

#### *CTPS Activity Assay*

Purified CTPS protein was incubated in the reaction buffer containing 100 mM NaCl, 30 mM Tris-HCl pH 7.5, 1mM ATP, 1mM UTP, 0.2 mM GTP, and 10 mM MgCl2 for 15 min at 37 °C. To initiate reaction, 10 mM prewarmed glutamine was added into the mixture. For inhibitor assay, 0.4 mM CTP and 2 mM NADH or adenine were added to the mixture. Absorption of a wavelength of 291 nm of each reaction mixture was measured with SpectraMax i3 as the indication for CTP production at individual time points(1).

#### *Phylogenetic construction*

112 sequences of CTPS of different species were acquired from the Swiss-Prot database(2), and MUSCLE(3) program was used for sequence alignment. Phylogenetic tree was built by Unweighted Pair Group Method (UPGMA)(4) using MEGA11(5). Poisson regression method was chosen for the modeling and pairwise deletion was applied for gaps treatment.

#### *Design and structure prediction of engineered protein*

ecCTPShelix12+ was designed by inserting helix 12 from hCTPS1 into ecCTPS after E399. The structure of ecCTPShelix12+ was predicted using ColabFold(6), with the protein sequence ”MTTNYIFVTGGVVSSLGKGIAAASLAAILEARGLNVTIMKLDPYI NVDPGTMSPIQHGEVFVTEDGAETDLDLGHYERFIRTKMSRRNNF TTGRIYSDVLRKERRGDYLGATVQVIPHITNAIKERVLEGGEGHD VVLVEIGGTVGDIESLPFLEAIRQMAVEIGREHTLFMHLTLVPYM AASGEVKTKPTQHSVKELLSIGIQPDILICRSDRAVPANERAKIALF CNVPEKAVISLKDVDSIYKIPGLLKSQGLDDYICKRFSLNCPEANL SEWEQVIFEEANPVSEVTIGMVGKYIELPDAYKSVIEALKHGGLK NRVSVNIKLIDSQDVETRGVEEEPVRYHEAWQILKGLDAILVPGG FGYRGVEGMITTARFARENNIPYLGICLGMQVALIDYARHVANM ENANSTEFVPDCKYPVVALITEWRDENGNVEVRSEKSDLGGTMR LGAQQCQLVDDSLVRQLYNAPTIVERHRHRYEVNNMLLKQIEDA GLRVAGRSGDDQLVEIIEVPNHPWFVACQFHPEFTSTPRDGHPLF AGFVKAASEFQKRQAKHHHHHH” serving as the input. Among the five prediction results, the model of rank2 exhibited the best performance and was selected for the subsequent analysis.

***Cryo-EM Grid Preparation and Data Collection***

For preparing the ecCTPS filament sample, 7 μM CTPS was incubated with 0.58 mM DON, 1 mM CTP, 0.5 mM NADH, and 10 mM MgCl2 for 30 min at 0 °C. For preparing the ecCTPS tetramer sample, 4.5 μM CTPS was incubated with 0.58 mM DON, 2 mM ATP, 2 mM UTP, 0.2 mM

GTP, and 10 mM $MgCl_2$ for 30 min at 0 °C. Samples were prepared with 300 holey golden film (M01Au300-R1.2/1.3) and FEI Vitrobot (4 °C temperature, multiple rounds of sample application and blotting before vitrification, 3.5 s blotting time, −1 blot force). Images were taken with a Gatan K3 summit camera on an FEI Titan Krios electron microscope operated at 300 kV. The magnification was 22,500 × in superresolution mode with the defocus rage -1.2 to -1.8 μm and a pixel size of 1.06 Å. The total dose was 50e−/Å2 subdivided into 40 frames at 2.8-s exposure using SerialEM.

***Image Processing***

We used MotionCor2 for alignment of the raw movie frames, and CTFFIND4 for CTF estimation. Ultimately, we selected 2760 images for further processing. By employing template-free particle picking in Relion, we obtained coordinates for 3,401,539 particles. Following 2D classification with bin2, we retained 1,934,043 particles for 3D classification. After 3D classification with C1 symmetry, we selected 1,300,114 particles for D2 symmetry 3D classification. Subsequently, we re-extracted particles and performed 3D refinement. Mask focusing refinement, CTF refinement, and Bayesian polishing were applied to improve the map resolution. After these procedures, we obtained a map with a resolution of 2.9 angstroms. Subsequent quality enhancement of the map was performed using ResolveEM within Phenix, resulting in a final resolution of 2.8 angstroms.

#### *Model Building and Refinement*

We employed Model PDB ID 5U3C as the initial model. Initially, a rigid body fitting was conducted using Phenix(7), followed by manual coordinate adjustments using Coot(8). Subsequently, real-space refinement was carried out once again using Phenix.

#### *Statistical analysis*

Results of CTPS activity assay were analyzed using GraphPad Prism 8(9) and were shown as means ± SD of three or more independent experiments. MUSCLE program was used for alignment of sequence of human CTPS1 (UniProtKB:P17812) and CTPS2 (UniProtKB:Q9NRF8), Schizosaccharomyces pombe CTPS (UniProtKB:O42644), drosophila CTPS (UniProtKB:Q9VUL1), E.coli CTPS (UniProtKB:P0A7E5), Mycobacterium tuberculosis CTPS (UniProtKB:P9WHK7), Thermus thermophilus CTPS (UniProtKB:Q5SIA8), and Sulfolobus solfataricus CTPS (UniProtKB:Q980S6).The result of sequence alignment was visualized by ESPript 3(10) which rendered sequence similarities and structure information taking crystal structure of tetrameric form of human CTPS1 (PDB EntryID:7MGZ) as reference.

### Supplemental Table S1. Cryo-EM data collection and model refinement.

| Table S1. Cryo-EM data collection and model refinement | | |
|---|---|---|
| Model | PDB ID 8I9O EMDB-35278 | |
| **Data collection** | | |
| EM equipment | Titan Krios | |
| Detector | K3 camera | |
| Magnification | 22,500x | |
| Voltage (kV) | 300 | |
| Electron exposure ((e–/Å$^2$)) | 50 | |
| Defocus range(μm) | -0.8 to -1.6 | |
| Pixel size(Å) | 0.53 | |
| Symmetry imposed | D2 | |
| Number of collected movies | 4204 | |
| Initial particle images (no.) | 3401539 | |
| Final particle images (no.) | 1300114 | |
| **Refinement** | | |
| Composition | | |
| Chains | 16 | |
| Atoms | 16632 (Hydrogens: 0) | |
| Residues | Protein: 2112 Nucleotide: 4 | |
| Water | 0.00 | |
| Ligands | CTP: 4 | |
| | MG: 4 | |
| Bonds (RMSD) | | |
| Length (Å) | 0.003 (0) | |
| Angles (°) | 0.575 (1) | |
| MolProbity score | 1.95 | |
| Clash score | 7.07 | |
| Ramachandran plot (%) | | |
| Outliers | 0.38 | |
| Allowed | 4.16 | |
| Favored | 95.46 | |
| Rama-Z (Ramachandran plot Z-score, RMSD) | | |
| whole (N = 2216) | 0.59 (0.18) | |
| helix (N = 888) | 1.87 (0.19) | |
| sheet (N = 372) | 0.53 (0.23) | |
| loop (N = 956) | -1.22 (0.20) | |
| Rotamer outliers (%) | 2.13 | |
| C-α outliers (%) | 0.00 | |
| Peptide plane (%) | | |
| Cis proline/general | 0.0/0.0 | |
| Twisted proline/general | 0.0/0.0 | |
| CaBLAM outliers (%) | 0.00 | |
| ADP (B-factors) | | |
| Iso/Aniso | 16632/0 | |
| min/max/mean | | |
| Protein | 0.00/87.95/38.12 | |
| Nucleotide | 25.62/41.41/35.92 | |
| Ligand | 0.00/42.82/19.82 | |
| Water | --- | |
| Occupancy | | |
| Mean | 1.00 | |
| occ = 1 (%) | 100.00 | |
| 0 < occ < 1 (%) | 0.00 | |
| occ > 1 (%) | 0.00 | |
| Data | | |
| Box | | |
| Lengths (Å) | 136.74,93.28,135.68 | |
| Angles (°) | 90,90,90 | |
| Supplied Resolution (Å) | 2.80 | |
| Resolution Estimates (Å) | Masked | Unmasked |
| d FSC (half maps; 0.143) | --- | --- |
| d 99 (full/half1/half2) | 2.7/---/--- | 2.7/---/--- |
| d model | 2.80 | 2.80 |
| d FSC model (0/0.143/0.5) | 2.5/2.6/2.9 | 2.5/2.6/2.8 |
| Map min/max/mean | -8.90/13.84/0.00 | |
| Model vs. Data | | |
| CC (mask) | 0.86 | |
| CC (box) | 0.73 | |
| CC (peaks) | 0.75 | |
| CC (volume) | 0.83 | |
| Mean CC for ligands | 0.87 | |

Supplemental figures S1-S8

**A**

**B**

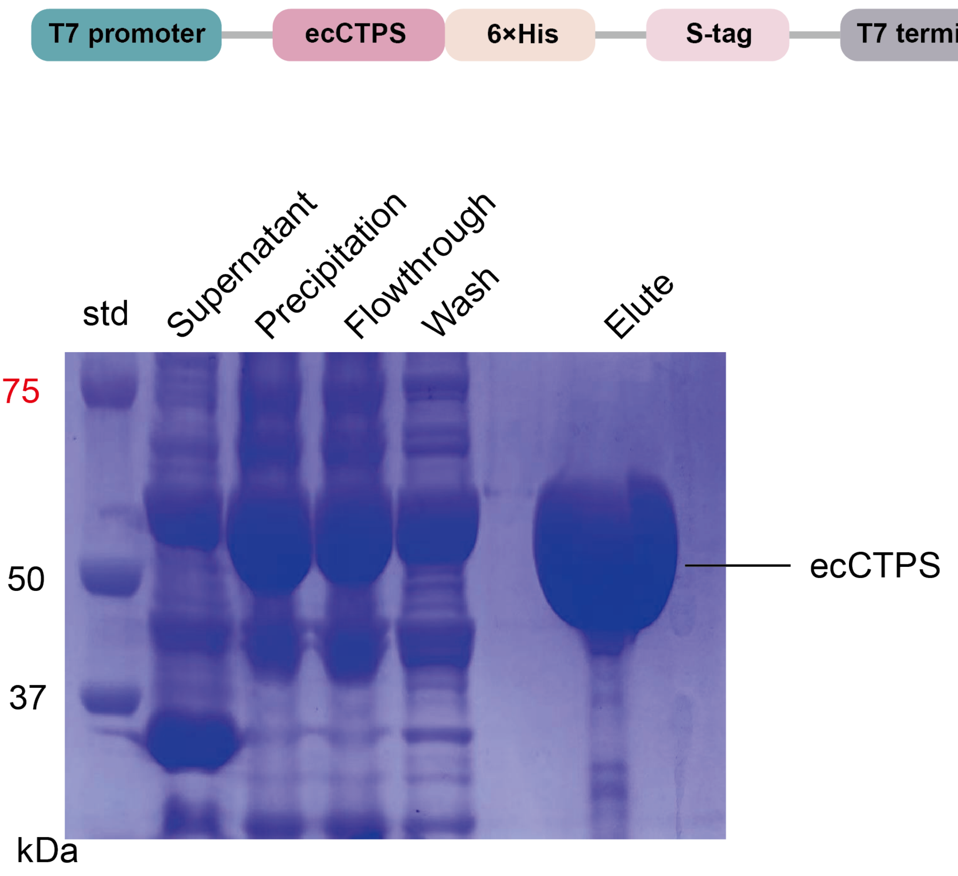


**Fig. S1. Construction, expression and purification of wild type ecCTPS protein.**
A) Plasmid construction. The plasmid was designed for the expression of ecCTPS with a 6×His tag at the C-terminus. B) SDS-Page analysis of purified ecCTPS proteins.

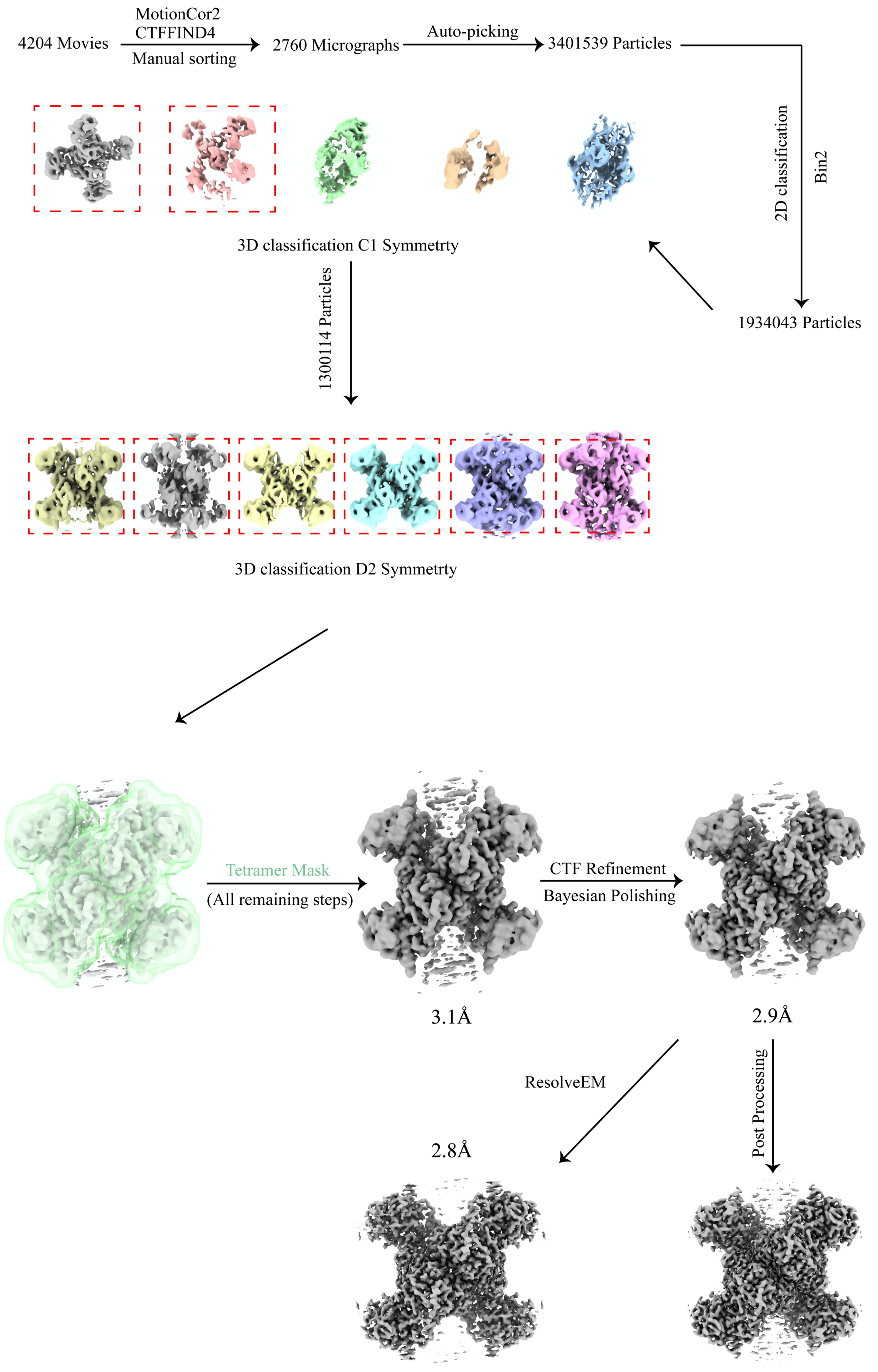


**Fig. S2. Work flow of Cryo-EM data processing.**

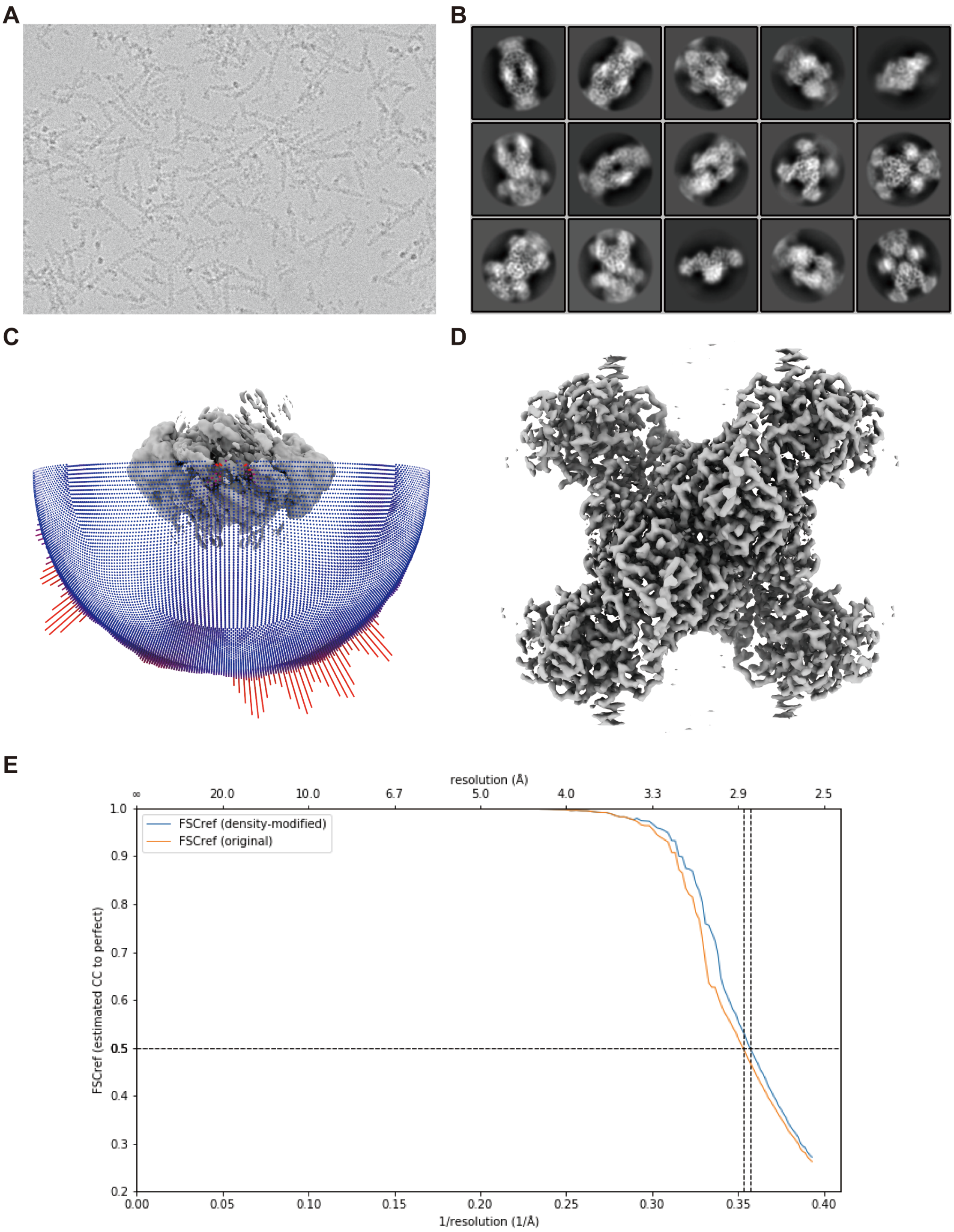


**Fig. S3 Properties of cryo sample and map.**

A) Image of cryo sample. B) Part of 2D classification result. C) Angular distribution of the final map. D) Map density from ResolveEM. E) FSC curve of the final map.

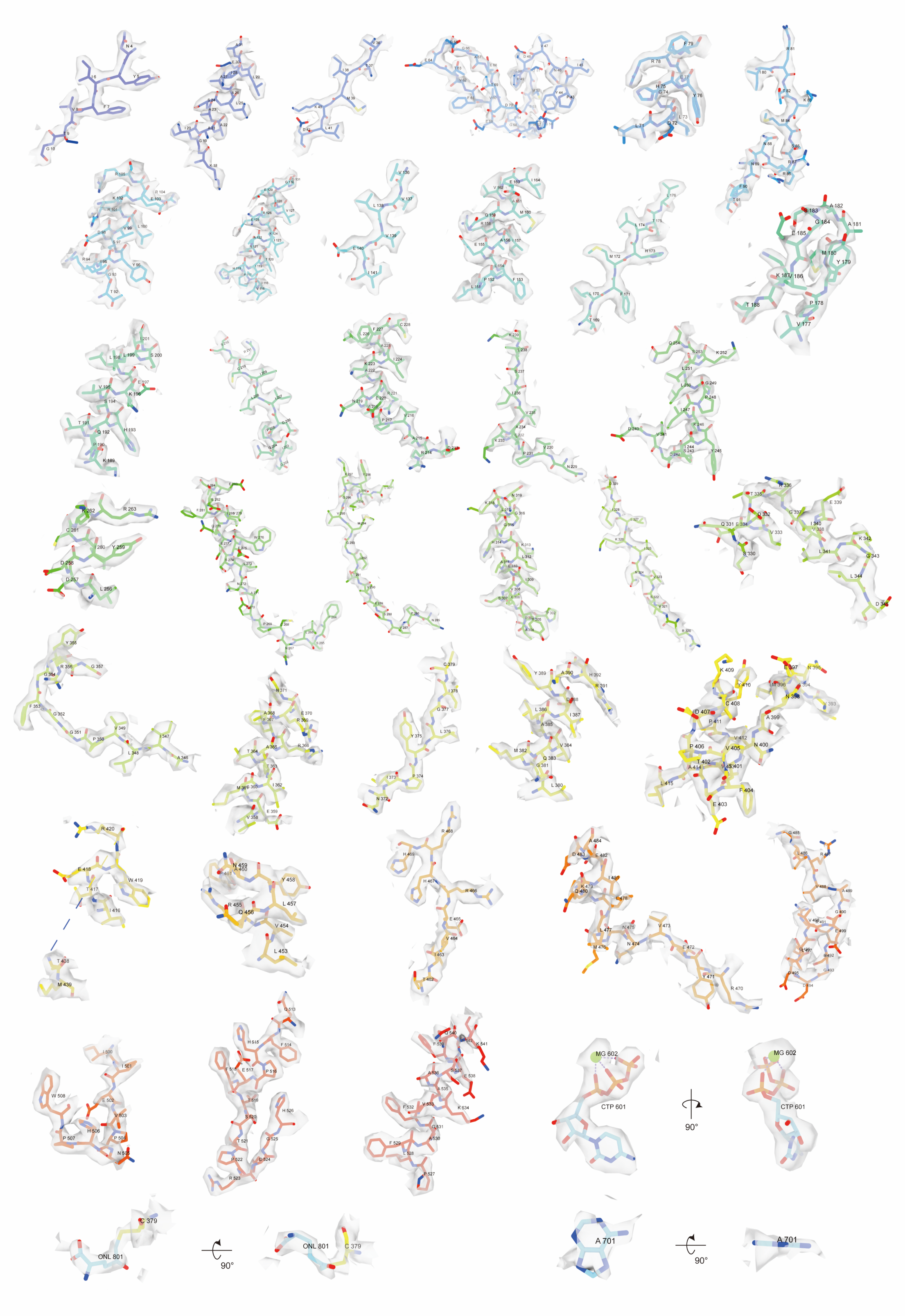

Fig. S4. Representative map density of model.

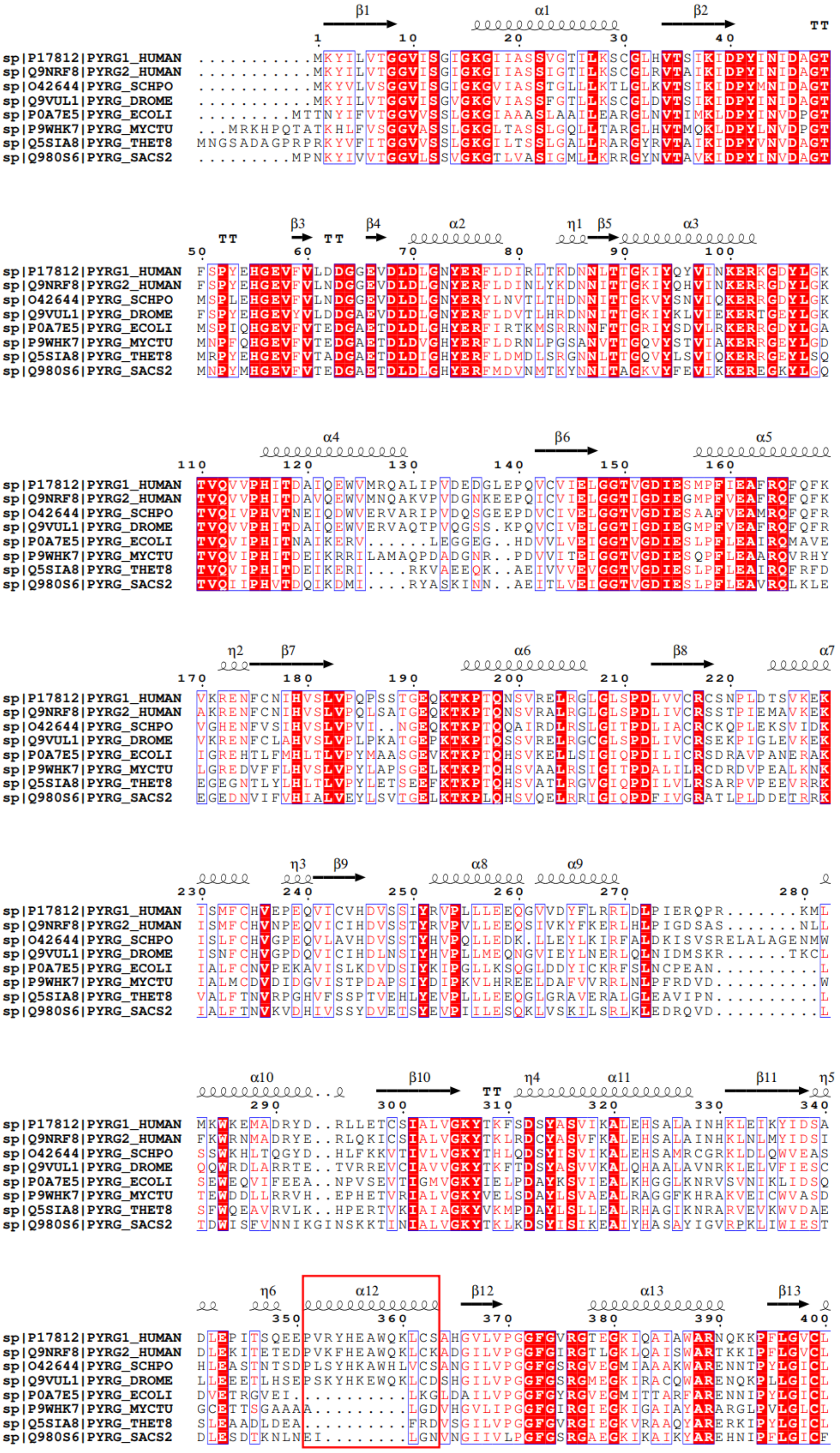
sp|P17812|PYRG1_HUMAN
sp|Q9NRF8|PYRG2_HUMAN
sp|O42644|PYRG_SCHPO
sp|Q9VUL1|PYRG_DROME
sp|P0A7E5|PYRG_ECOLI
sp|P9WHK7|PYRG_MYCTU
sp|Q5SIA8|PYRG_THET8
sp|Q980S6|PYRG_SACS2

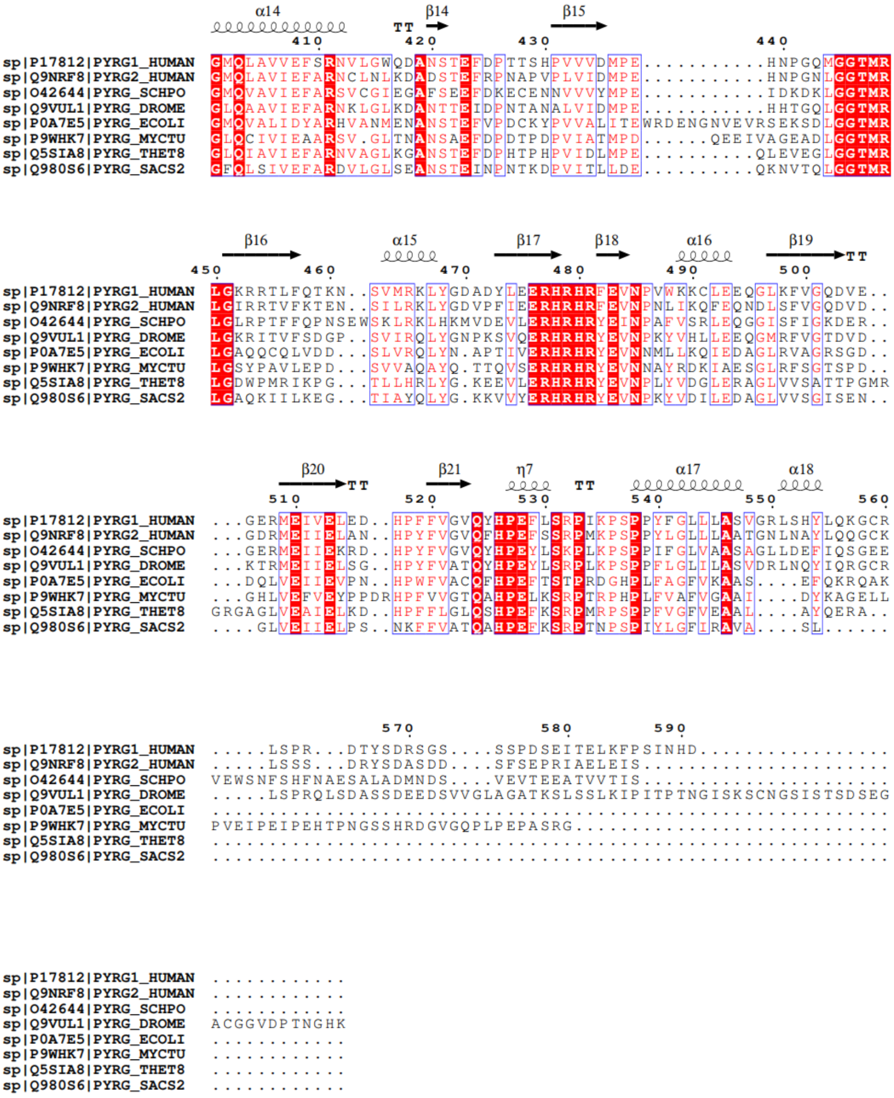

**Fig. S5. Seqence alignment of CTPS from different species.**

The sequence alignment of the amino acid sequence of human CTPS1 and CTPS2, Schizosaccharomyces pombe CTPS, drosophila CTPS, e.coli CTPS, Mycobacterium tuberculosis CTPS, Thermus thermophilus CTPS, and Sulfolobus solfataricus CTPS is shown. The conserved residues are shaded in red, and secondary structure information is shown above the alignment. The red box indicates the deletion of alpha helix 12 in archaeas and prokaryotes.

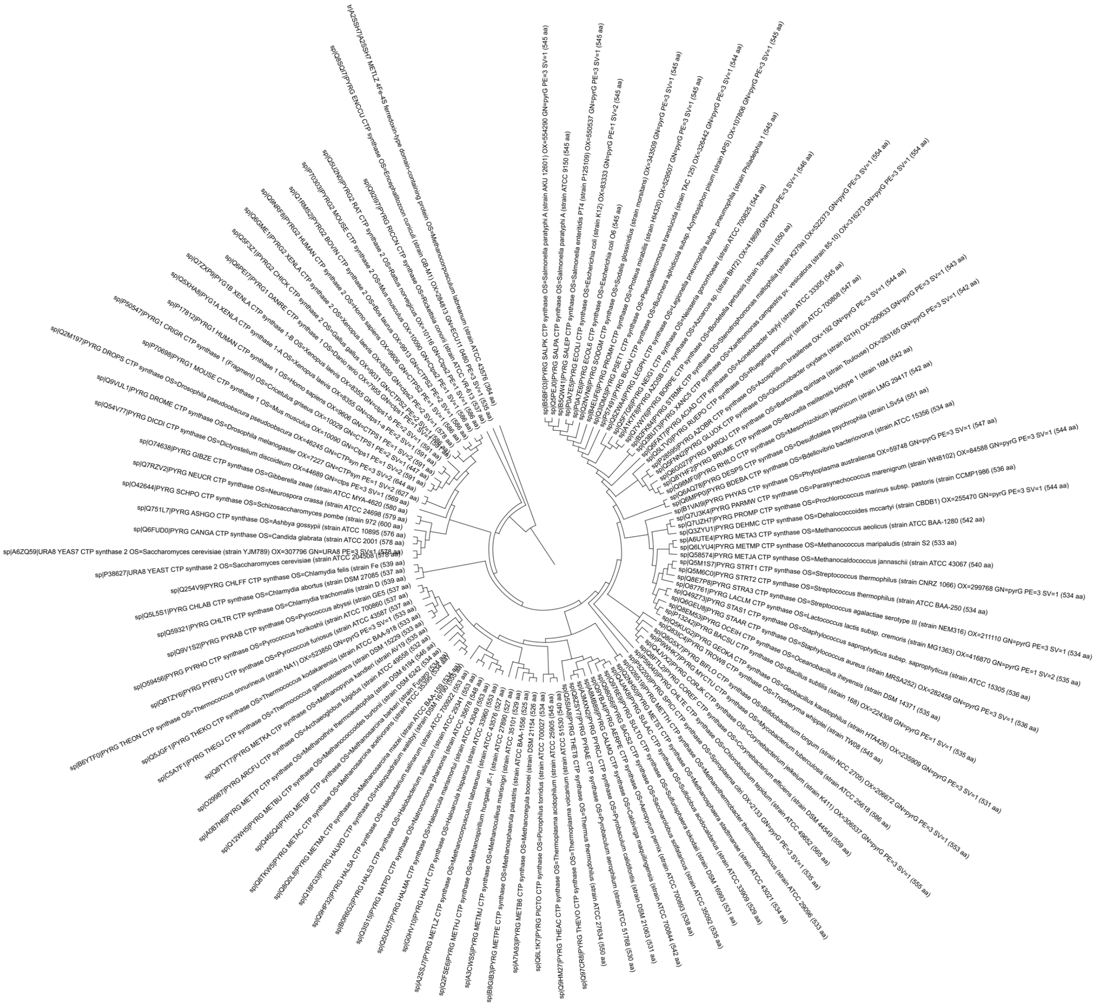

**Fig. S6. Phylogenetic tree.**

112 sequences of CTPS of different species were acquired from the Swiss-Prot database and used to build the phylogenetic tree.

A

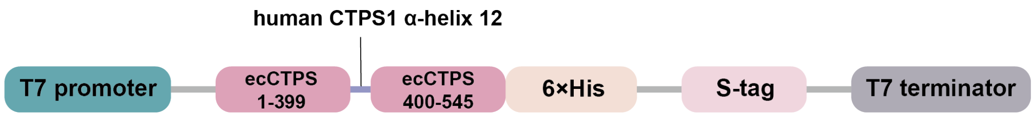


B

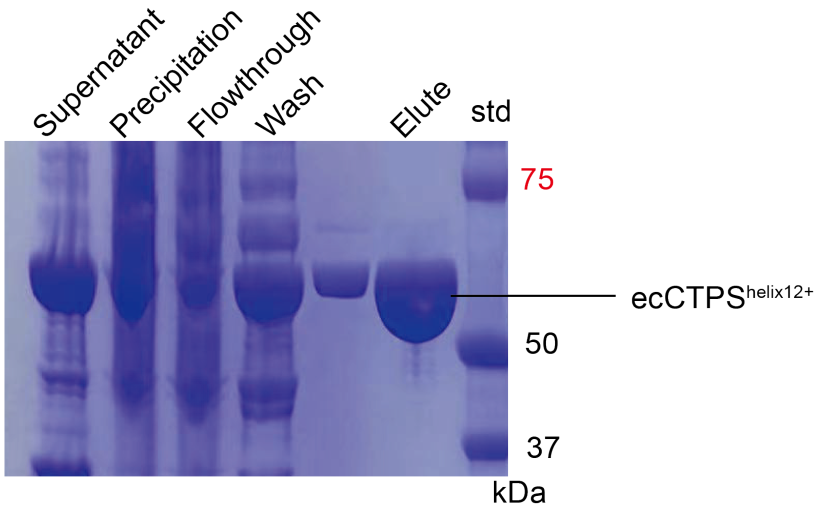


**Fig. S7. Construction, expression and purification of ecCTPS$^{helix12+}$ protein.**

A) Plasmid construction of ecCTPS$^{helix12+}$ protein. Human CTPS1 alpha helix 12 (EEPVRYHEAWQ) was inserted into ecCTPS after the 399th amino acid. The plasmid was designed for the expression of ecCTPS$^{helix12+}$ with a 6×His tag at the C-terminus. B) SDS-Page analysis of purified ecCTPS$^{helix12+}$ proteins.

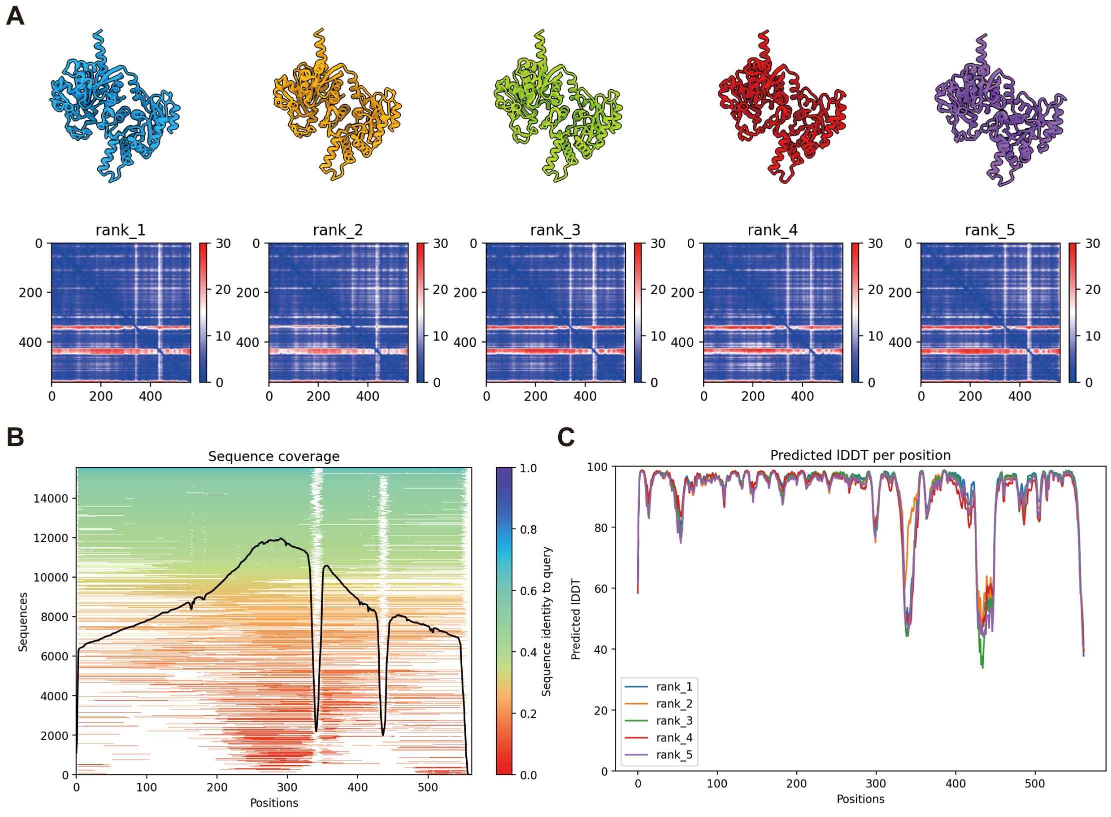


**Fig. S8. Structure prediction of ecCTPS$^{helix12+}$ by AlphaFold2.**

A) Models and their inter-chain predicted alignment error (inter-PAE) provided by AlphaFold2. B) Visualization of multiple sequence alignment depth and diversity. C) Plots of predicted local distance difference test(pLDDT) per models.
